# Genome Editing in *Caenorhabditis briggsae* using the CRISPR/Cas9 System

**DOI:** 10.1101/021121

**Authors:** Elizabeth Culp, Cory Richman, Devika Sharanya, Bhagwati P Gupta

**Affiliations:** Department of Biology, McMaster University, 1280 Main Street West, Hamilton, ON L8S-4K1, Canada

**Keywords:** CRISPR/Cas9, genome editing, nematode, *C. briggsae*

## Abstract

**Author Summary:** The CRISPR/Cas9 system has recently emerged as a powerful tool to engineer the genome of an organism. The system is adopted from bacteria where it confers immunity against invading foreign DNA. This work reports the first successful use of the CRISPR/Cas9 system in *C. briggsae*, a cousin of the well-known nematode *C. elegans*. We used two plasmids, one expressing Cas9 endonuclease and the other an engineered CRISPR RNA corresponding to the DNA sequence to be cleaved. Our approach allows for the generation of loss-of-function mutations in *C. briggsae* genes thereby facilitating a comparative study of gene function between nematodes.

**Abstract:** The CRISPR/Cas9 system is an efficient technique for generating targeted alterations in an organism’s genome. Here we describe a methodology for using the CRISPR/Cas9 system to generate mutations via non-homologous end joining in the nematode *Caenorhabditis briggsae,* a sister species of *C. elegans*. Evidence for somatic mutations and off-target mutations are also reported. The use of the CRISPR/Cas9 system in *C. briggsae* will greatly facilitate comparative studies to *C. elegans.*

---

Linking genotype and phenotype is an important step in the characterization of a gene. Targeted genome editing, defined as the creation of alterations at specific sites in an organism’s genome, is a powerful means to study the relationship between gene and phenotype. Genome editing techniques are based on guiding an endonuclease to a specific target in the genome in order to generate a double strand break (DSB) [1–3]. Breaks are subsequently repaired by either error prone non-homologous end joining (NHEJ) or template-directed homologous recombination (HR) [4]. While the former introduces random mutations at the point of cleavage, the latter can be used to generate specific alterations based on the presence of a donor sequence. Although several technologies currently exist for genome editing, such as zinc finger nucleases (ZFN) and transcription activator-like effector nucleases (TALEN), these techniques leave room for improvement in their ease of use, as each new sequence to be targeted requires the labor intensive process of generating a new protein construct [2].

Clustered, regularly interspaced, short palindromic repeats (CRISPR) and CRISPR- associated (Cas) systems are adaptive immune mechanisms evolved by archaea and bacteria to defend against foreign plasmids and viral DNA [5]. Manipulation of the *Streptococcus pyrogenes* type II CRISPR/Cas system has been used to develop an efficient genome editing technique. First, a 20 bp sequence in a gene of interest is selected to act as a guide for the *S. pyrogenes* nuclease, Cas9. This sequence, termed the CRISPR RNA (crRNA), has the only requirement that it must precede a Protospacer Adjacent Motif (PAM) of the form 3’NGG [6]. Next, a second RNA molecule, termed the trans-activating crRNA (tracrRNA), is used for binding to Cas9 [6]. For the purpose of experimental simplification, the crRNA and tracrRNA sequences can be fused into a single guide RNA (sgRNA) [7]. By expressing this sgRNA along with Cas9 in germ line cells, heritable genome mutations can be created.

The CRISPR/Cas9 system has been successfully established in two leading nematode models – *C. elegans* and *Pristionchus pacificus* [2, 8]. Friedland *et al.* [9] developed a simple protocol for *C. elegans* that involved injecting plasmids into the gonad of adult hermaphrodites. The authors modified Cas9 to include a SV40 NLS to ensure nuclear localization and expressed under an *eft-3* translation elongation factor promoter, chosen for its effectiveness in germ line expression. The sgRNAs were expressed under a U6 small nuclear RNA polymerase III promoter, chosen for its ability to drive expression of small RNAs. As the optimal expression from this promoter requires the first base to be a purine, the sgRNA target sequence is restricted to the form (G/A)(N)_19_NGG [9, 10].

Adaptation of CRISPR/Cas9 to *C. briggsae,* a species that is closely related to *C. elegans,* would provide a powerful tool to investigate the function of any given gene. *C. briggsae* is used routinely by many laboratories in comparative evolutionary studies. The two animals diverged less than 30 million years ago yet share similar morphology [11]. A comparison of their genome sequences has revealed that roughly one-quarter of their genes lack clear orthologs including many that are highly divergent and species-specific [12]. This suggests that underlying gene networks have evolved substantially without an obvious change in phenotype [13]. Such changes are likely to have significant impacts and may confer unique advantages on animals to withstand genetic and environmental fluctuations. By generating mutations in *C. briggsae* genes and characterizing phenotypes, we can learn the functional relevance of genomic differences, including any alterations in genetic pathways and developmental mechanisms between the two species. With this goal in mind, we set out to develop a method for using this system in *C. briggsae*.

The wild type AF16 strain was used as a reference strain in all experiments. Strains generated as part of this study include DY503 *Cbr-unc-22(bh29)*, DY504 *Cbr-dpy-1(bh30),* DY530 *Cbr-bar-1(bh31),* DY544 *Cbr-unc-119(bh34)* and DY545 *Cbr-unc-119(bh35).*

We first used the CRISPR/Cas9 system in *C. briggsae* in an attempt to generate targeted loss-of-function mutations by employing NHEJ. For this, two conserved genes were chosen based on visible phenotypes, *Cbr-dpy-1,* a cuticle protein causing a dumpy (Dpy) phenotype, and *Cbr-unc-22,* a twitchin homolog causing an uncoordinated (Unc) phenotype [14–16]. Target sgRNA sequences following the form G/A(N)_19_NGG were searched for in the exonic regions of these genes using the ZiFiT Targeter Version 4.2 software [17]. The sgRNA sites were screened based on predicted efficiency using empirically based scoring algorithms. Off-target sites were minimized using the sgRNAcas9 software package developed by Xie *et al.* [18].

The plasmids containing the *C. elegans* U6 promoter and sgRNA target sequences were generated by site-directed mutagenesis. This was accomplished using either two-step overlap-extension PCR on a *pU6::Cbr-unc-119_sgRNA* template (gift from John Calarco, Addgene plasmid #46169) [9], or Q5 site-directed mutagenesis on a *pU6::Cbr-lin-10_sgRNA* template [19] using the NEB Q5 site-directed mutagenesis kit (E0554). The target site substitution was confirmed by *AclI* digestion. See Tables S1 and S2 for sgRNA sites and primers used in this study.

The plasmids sgRNA and Cas9 (*Peft-3::Cas9-SV40 NLS::tbb-2 3’UTR*, also from John Calarco, Addgene #46168) were injected into the germline of young adults using standard methods [20] and F1 progeny displaying the co-injection marker, pharyngeal expression of GFP, were isolated onto separate plates. Injection mixes contained *pU6::sgRNA* (100 ng/ul), *Peft-3::Cas9-SV40 NLS::tbb-2 3’UTR* (100 ng/ul), and *myo-2::GFP* (10 ng/ul).

Following microinjection, F2 worms were screened for desired phenotypes. We successfully isolated mutants for both *Cbr-dpy-1* and *Cbr-unc-22* at comparable frequencies to those observed in *C. elegans* (Table 1) [9]. Sequencing of the alleles of each of these genes revealed insertions and deletions at the sgRNA target sites (Table 2). The phenotypes of mutant animals are indistinguishable from those in *C. elegans* corresponding to orthologous genes, demonstrating conservation of gene function. Together, these results show that the CRISPR/Cas9 system works in *C. briggsae* and can utilize conserved *C. elegans* promoters to express sgRNAs and Cas9.

**Table 1.**
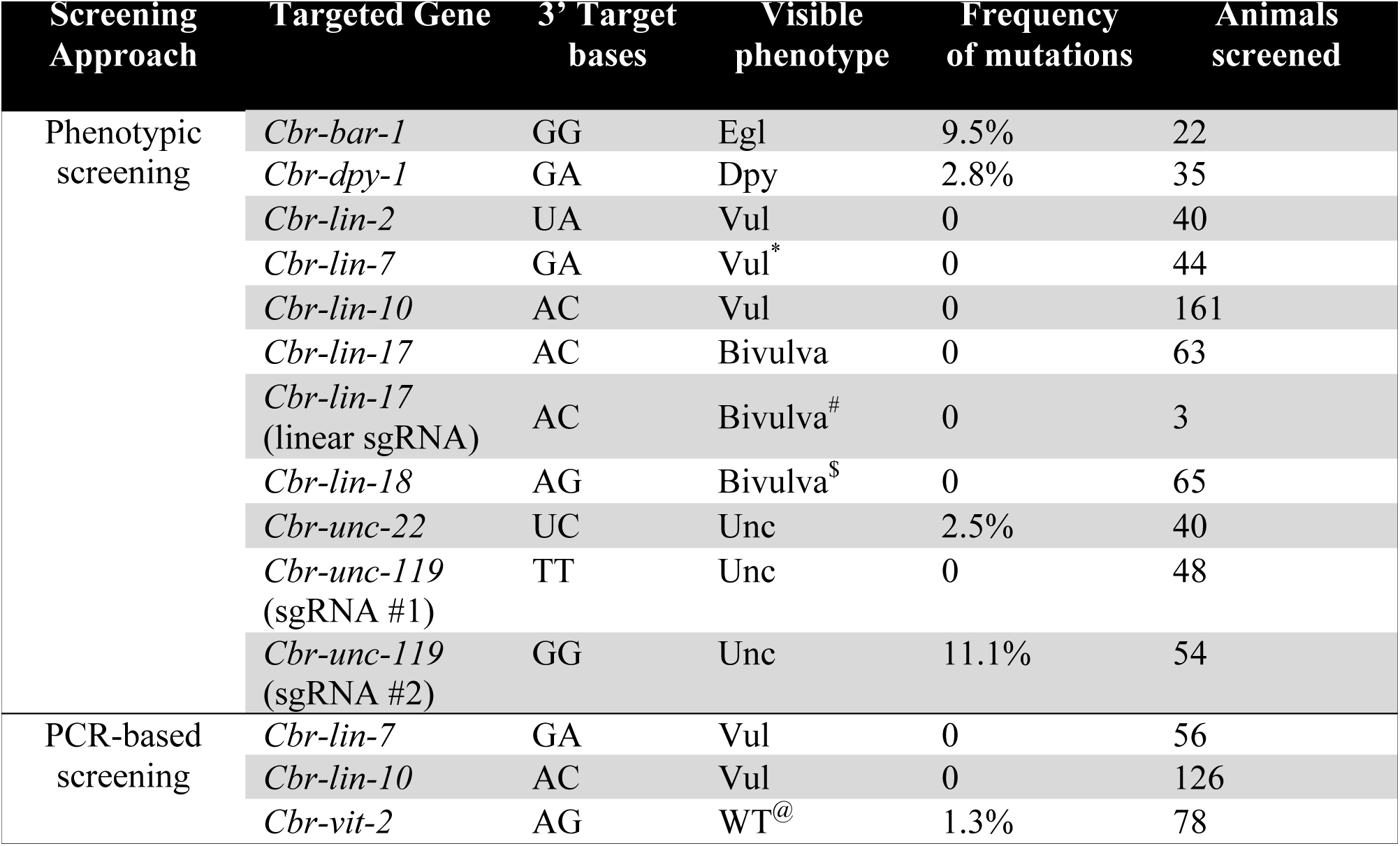
Phenotypes of transgenic animals generated using the CRISPR/Cas9 technique. The 3’ target bases are those at positions 19 and 20 in the sgRNA target sequence. ^*^One F2 showed Dpy phenotype. ^#^3 bivulva worms were recovered in F3 but the phenotype was not heritable. ^$^One F2 showed protruding vulva (Pvl) phenotype. ^@^wild type based on the *C. elegans vit-2* mutant phenotype.

**Table 2.**
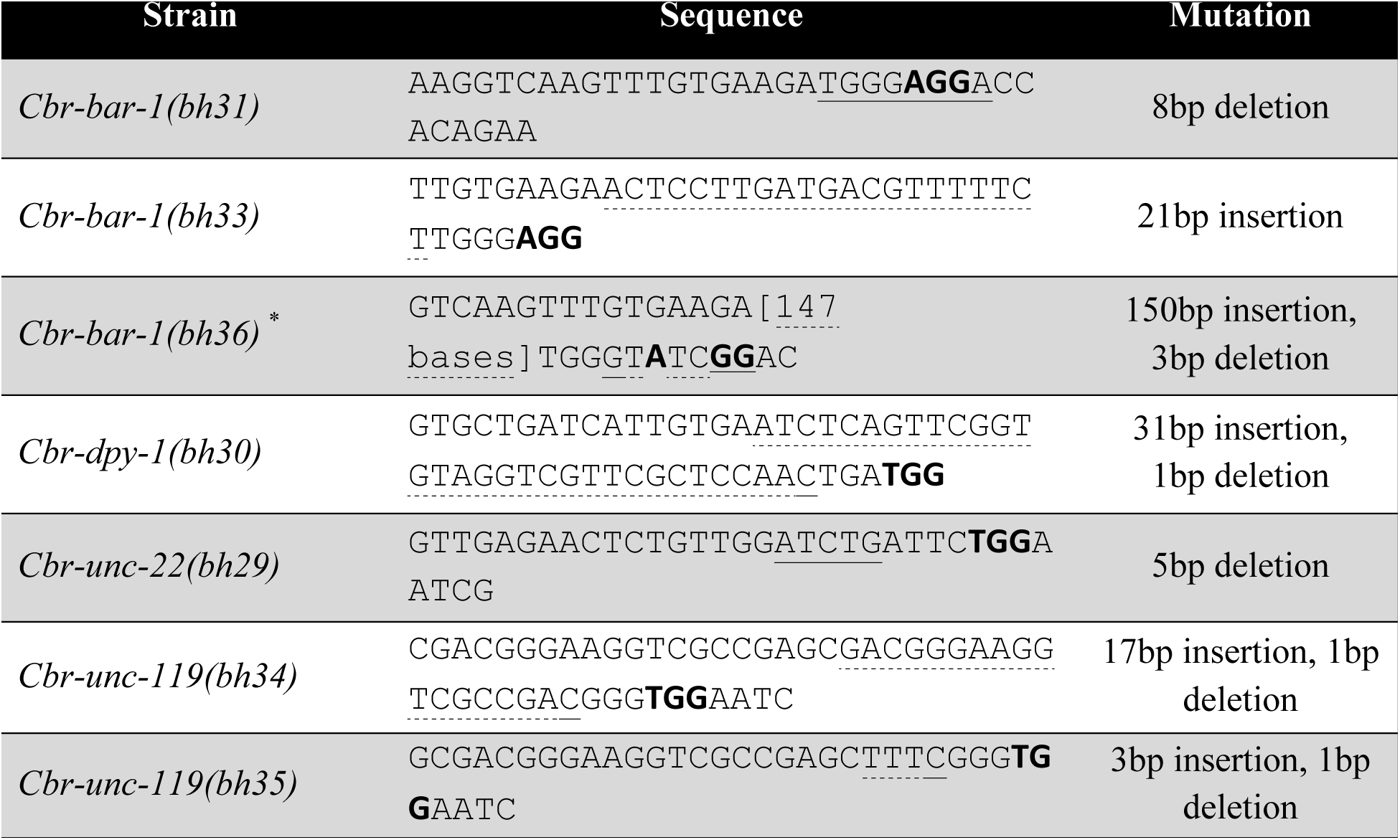
Alleles generated by the CRISPR/Cas9 approach. The DNA sequence includes the sgRNA target. The PAM site is bolded. Insertion and deletion sequences are underlined (dotted underline: insertion, solid underline: deletion). For clarity the 147 base pair inserted sequence in *bh36* allele has been omitted. This long sequence matches with the *E. coli* gene EF-Tu. ^*^The allele was recovered in a separate screen along with another allele *bh32* that has small deletion. The exact base change in *bh32* has not been determined.

Next, we targeted six other conserved genes of the Wnt and Ras pathways (*Cbr-lin-2, Cbr-lin-7, Cbr-lin-10, Cbr-lin-17, Cbr-lin-18* and *Cbr-vit-2*). For the PCR-based assay [19] F1s were allowed to lay eggs for 24-36 hours, and then picked and lysed in pools of two. A region of the genomic DNA spanning the sgRNA site (∼200 bp) was amplified and examined on a 4% high-resolution agarose gel (Invitrogen UltraPure Agarose-1000, Catalog #16550-100) for changes in band sizes (Figure S2). In some cases we recovered mutations as determined by phenotypic as well as PCR-based screening approaches but none were found to be heritable (Table 1). It is unclear to us whether it was due to sgRNAs being non-functional, less efficient or requiring much larger F1s to be screened. Similar results were previously reported in *C. elegans* [21]. In one case, *Cbr-lin-17*, we sequenced the animal that showed bi-vulva phenotype and found possible evidence for a somatic mutation (T/A transversion causing M482L substitution).

The bi-vulva phenotype in this line was lost in subsequent generations. Evidence of somatic mutations has also been described in *C. elegans* [21].

Interestingly, our screens also recovered worms with unexpected phenotypes, e.g., Dpy in *Cbr-lin-7* screen (Table 1). Sequencing of these worms revealed no disruption in targeted genes, raising the possibility of off-target effects of CRISPR/Cas9. Off target effects have been reported in *C. elegans* as well as several other models including *Drosophila,* mice, zebrafish, and human cell lines [22–25].

The sgRNAs with a 3’GG motif at positions 19 and 20 were recently shown to significantly enhance the efficiency of targeted mutations in *C. elegans* [21](23). To test whether a similar sequence structure could be effective in *C. briggsae* we selected two conserved genes *Cbr-unc-119* and *Cbr-bar-1*. Mutations in *Cbr-unc-119* with Unc phenotype were recovered at a frequency of 11.1% (Tables 1 and 2). In contrast, another sgRNA for *Cbr-unc-119* that lacked 3’ GG motif did not give rise to any mutation (Table 1). In the case of *Cbr-bar-1,* a β-catenin homolog [26], the 3’GG motif sgRNA resulted in a disruption efficiency of 9.5% (Tables 1 and 2). The enhanced efficiency of the 3’GG motif sgRNA sites for these two genes suggests that such an approach in *C. briggsae* could improve the frequency of targeted mutations in genes of interest.

In addition to the CRISPR-mediated NHEJ approach we also attempted the HR method of gene editing in *C. briggsae.* For this donor templates were designed to either disrupt a gene (by inserting a single-stranded oligonucleotide) or tag genes using double-stranded linear PCR amplicons (or plasmids) of fluorescent reporters (GFP and dsRED). Specifically, the single strand oligonucleotide donor templates were intended to insert a 22 bp sequence containing an *NcoI* restriction enzyme site into *Cbr-bar-1* and *Cbr-lin-15B* (Figure S1B). Homology arms of length 75 and 49 bases were chosen directly overlapping the sgRNA site, based on previous results [19]. The double-stranded linear donor templates of *GFP* (864 bp) and *dsRED* (830 bp) containing short microhomology arms were generated by PCR to create translational fusions with *Cbr-bar-1* and *Cbr-vit-2,* respectively (Figure S1C). The donor vector *myo-2::dsRED::unc-54 3’UTR* was designed to insert a *myo-2::dsRED* reporter into the *Cbr-bar-1* (Figure S1A) [27]. The vector contained a 2 kb transgene flanked on either side by 1 kb of sequence homologous to *Cbr-bar-1* (Gibson Assembly Cloning Kit NEB catalog #E5510). The templates were included in the injection mix (donor plasmid 200 ng/μl, linear PCR amplicons 50 ng/μl, single-stranded oligonucleotides 30 ng/μl) along with other DNA components as mentioned above. Although none of these HR approaches were successful, in some cases we did observe expected genomic changes in F1 and F2 animals (as determined by sequencing), which were not inherited in subsequent generations (Table 3).

**Table 3.** Genome editing events detected using CRISPR-mediated HR. The sgRNA efficiency shows all genome editing events, including those repaired by NHEJ and HR, based on phenotypic and PCR-based screens. HR efficiency indicates the number of HR events detected in F2 out of the total F1s screened. ^#^Wild type based on the phenotype of *C. elegans* orthologs.

| Targeted Gene | Expected phenotype | sgRNA Efficiency | HR Efficiency |
| --- | --- | --- | --- |
| *Cbr-bar-1* | Egl | 25/219 (11.4%) | 0/219 |
| *Cbr-bar-1* | Egl | 18/211 (8.5%) | 0/211 |
| *Cbr-bar-1* | Egl | Not Determined | 0/202 |
| *Cbr-lin-15B* | WT<sup>#</sup> | Not Determined | 0/68 |
| *Cbr-vit-2* | WT<sup>#</sup> | 1/78 (1.3%) | 0/78 |

In conclusion, we have shown that the CRISPR/Cas9 system can be effectively employed in *C. briggsae* to alter a gene of interest. Similar to *C. elegans* the 3’ GG motif appears to increase the frequency of NHEJ events. Interestingly, we observed a significant bias towards insertion NHEJ events in *C. briggsae.* Of the total of 8 alleles recovered, for 4 different genes, 62% had insertion of bases of varying length (range 3 to 150). Similar screens in *C. elegans* have reported 26% frequency of such events (n = 86 from 5 different studies) [9, 21, 28–30]. More work is needed to ascertain if such a bias in *C. briggsae* holds true in a larger sample size.

Together with the recently developed TALEN-based genome editing approach [3], the CRISPR/Cas9 approach described here provides a powerful means to investigate the functions of conserved as well as divergent genes in *C. briggsae*. This promises to accelerate comparative studies with *C. elegans* thereby leading to a greater understanding of the flexibility of genetic and molecular mechanisms during animal development.

## Acknowledgements

We thank all members of the Gupta lab, particularly Scott Amon, Ayush Ranawade and Anand Adhikari, for their input and assistance throughout this project. We also thank John Calarco for two plasmids and initial advice in microinjection experiments. This work was supported by funds from the Natural Sciences and Engineering Research Council of Canada (NSERC) Discovery program to BPG.

## Supplementary Materials

**Table S1.**
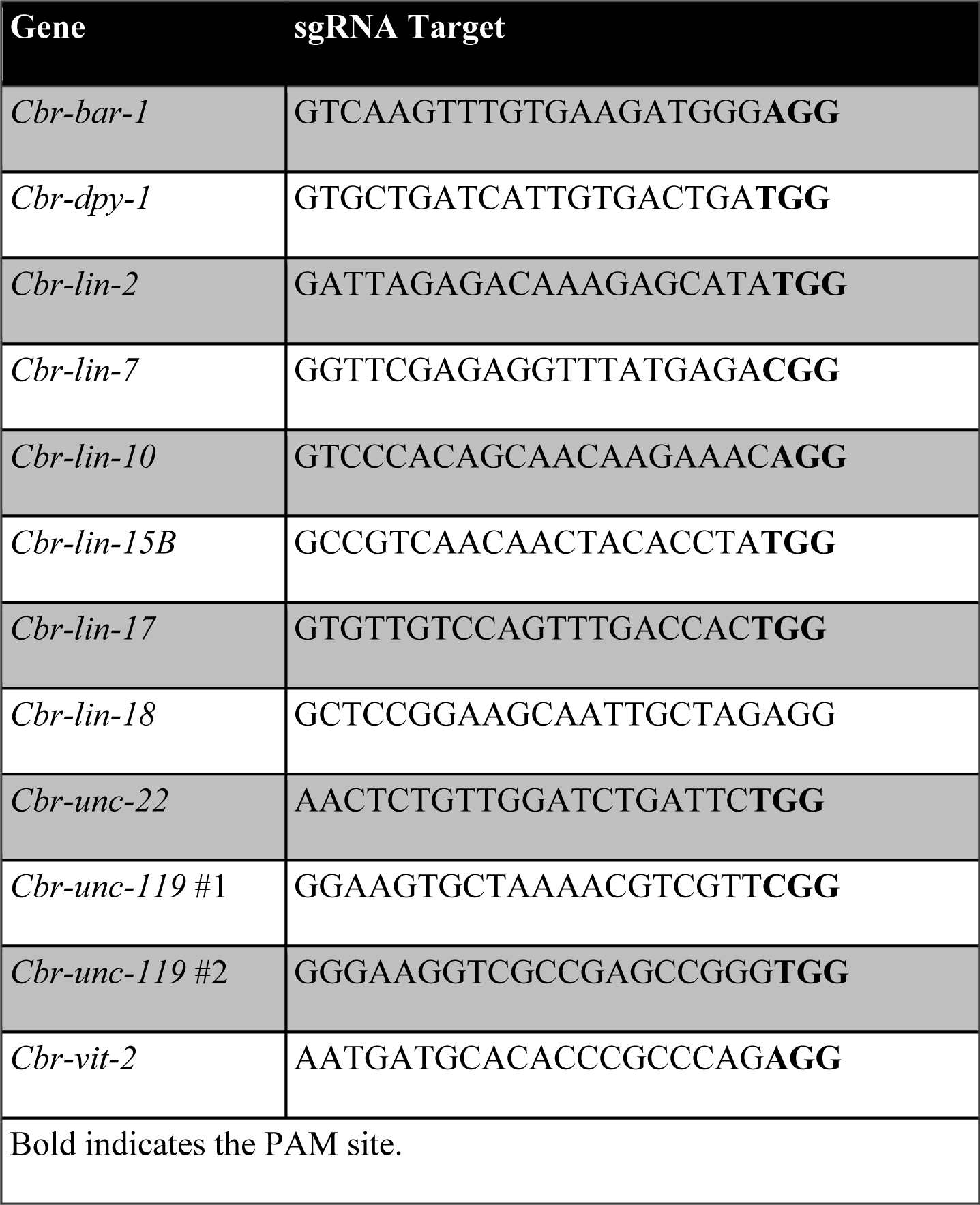
sgRNA target sites.

**Table S2.**
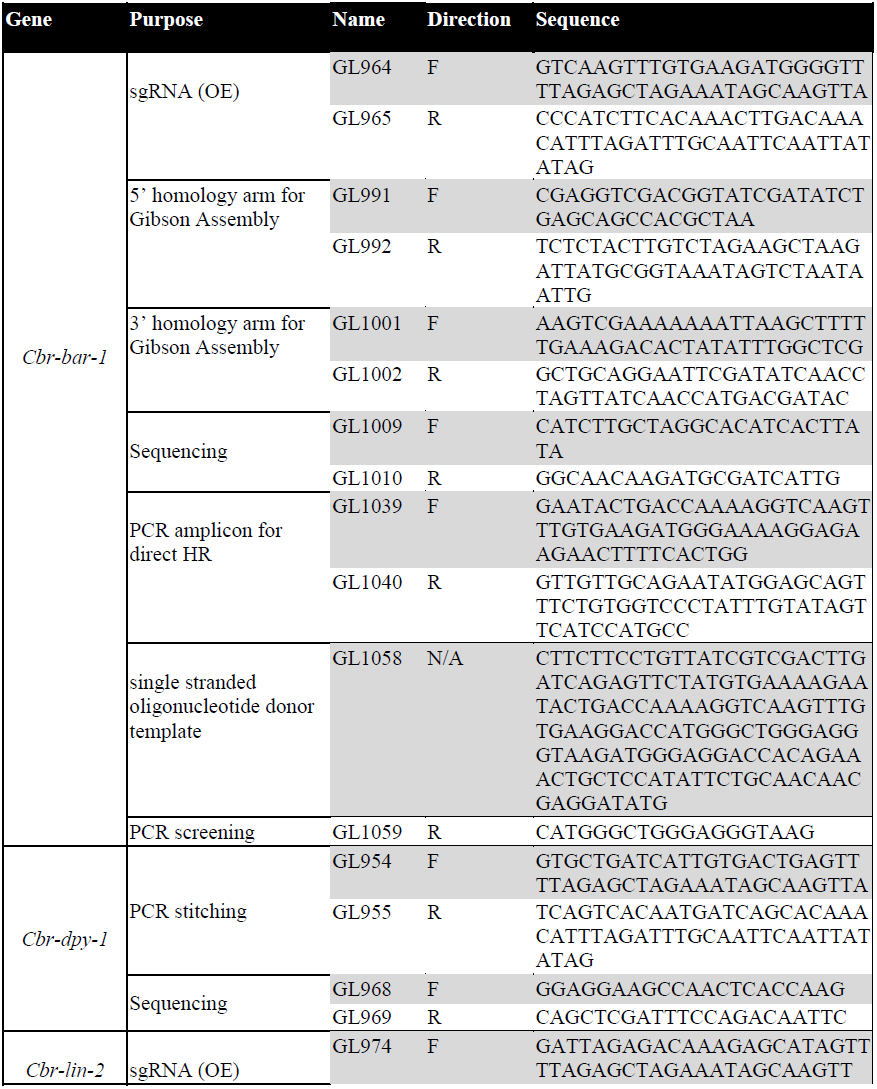
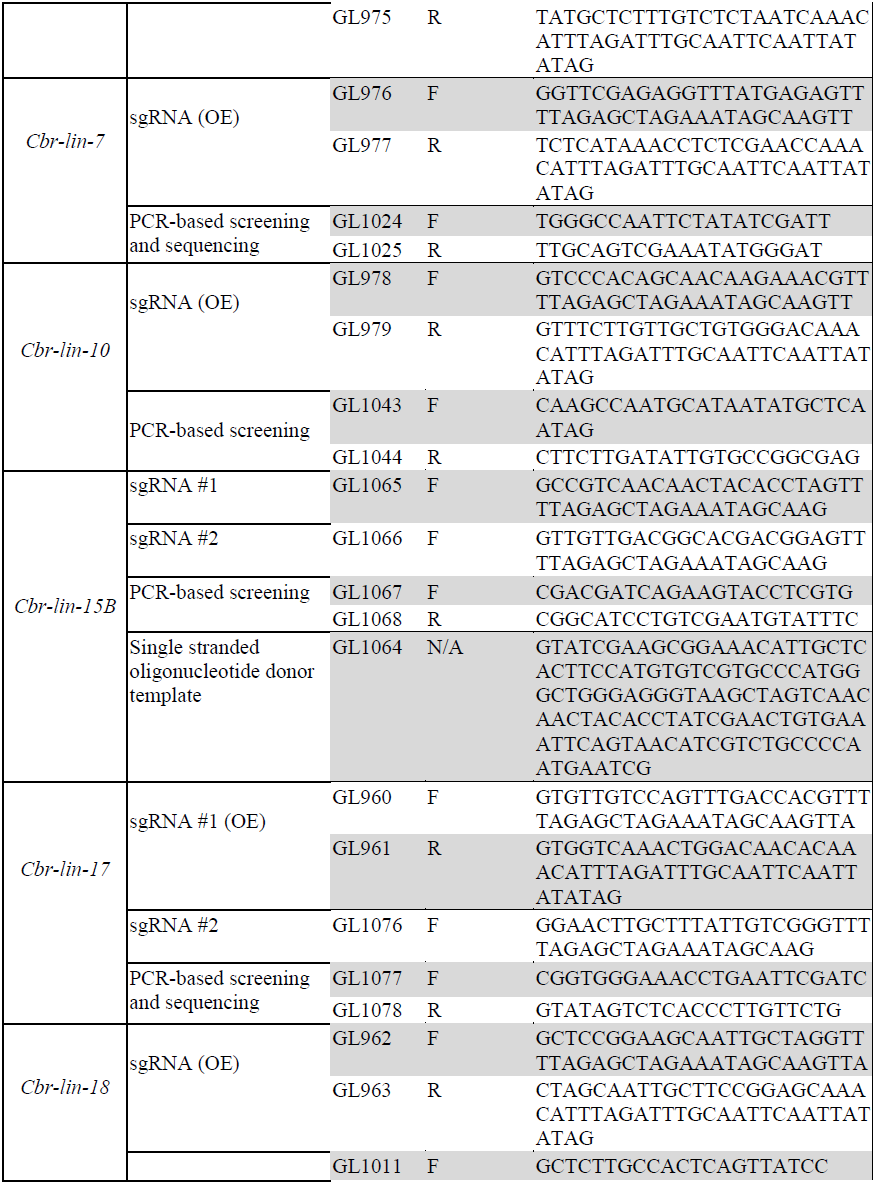
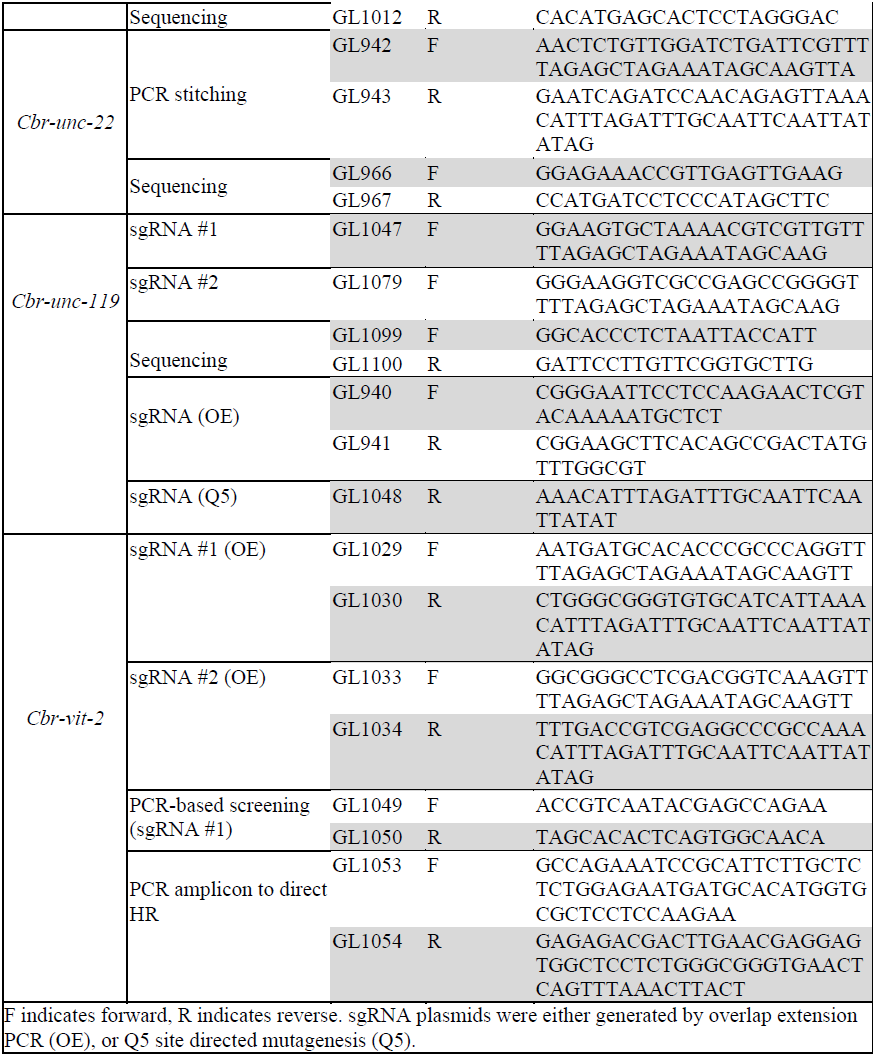
List of Primers.

**Figure S1.**
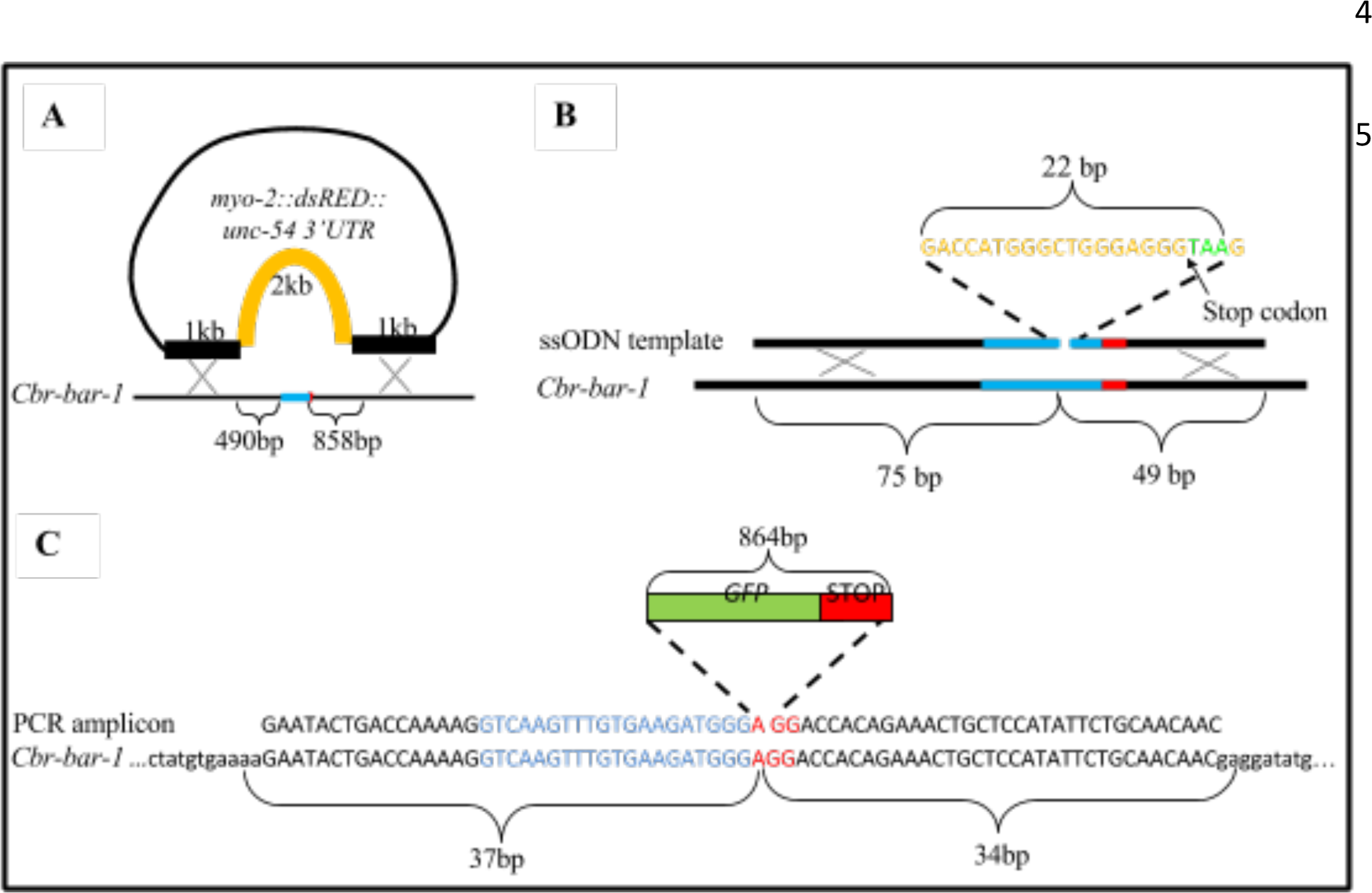
Donor sequence approaches generated as templates for HR for Cbr-bar-1.

Templates may take the form of a donor vector (A), ssODN (B) or PCR amplicons (C). Blue letters represent the sgRNA target sequence while red letters represent the PAM site.

**Figure S2.**
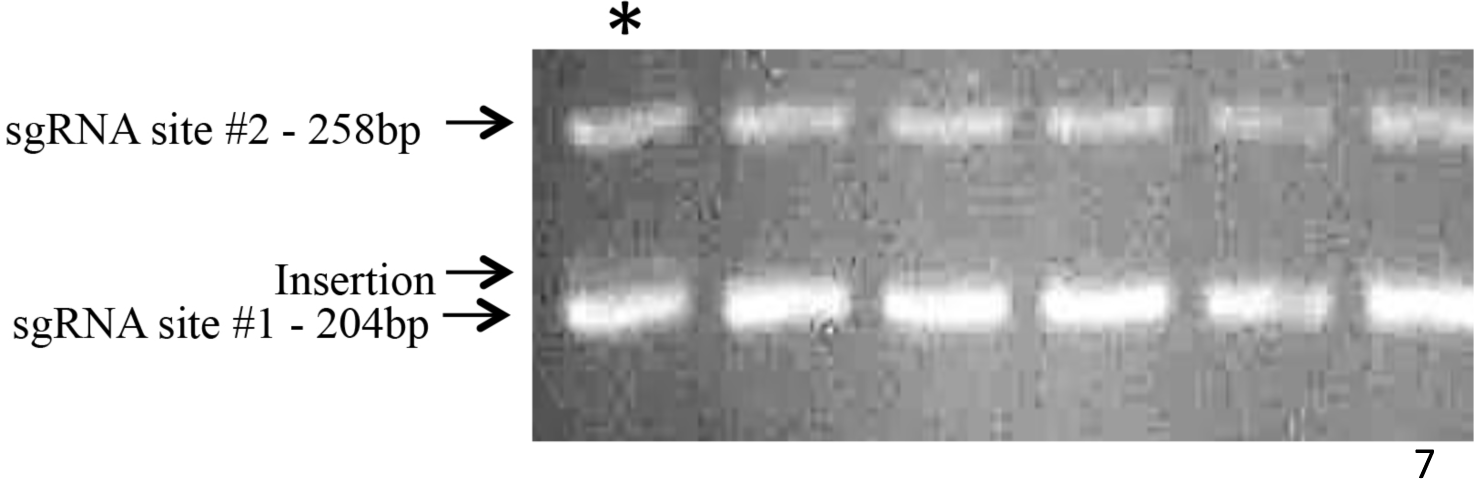
PCR amplicons of the *Cbr-vit-2* genomic region flanking the sgRNA target site.

An insertion can be seen at sgRNA site #1 in the lane marked with *.

